# Longitudinal Changes in Social–Alcohol Behavioral Allocation During Chronic Ethanol Exposure

**DOI:** 10.64898/2026.11.24.768927

**Authors:** Nathan Jones, Elliot Arozqueta, Owen Alt, Hannah Aranda, Phillip Starski, Doo-Sup Choi

## Abstract

**Background:** Conventional measures of ethanol intake do not capture how behavior is allocated between alcohol-associated and competing social options, nor how this allocation changes within the same individual over time. A modified three-chamber Social Alcohol Choice Test (SACT) was implemented to quantify within-individual changes before and after chronic alcohol exposure. Exploratory baseline stratification using the Chamber Sociability Index (CSI) enabled detection of heterogeneity in longitudinal response patterns.

**Methods:** Fifteen male C57BL/6J mice underwent baseline SACT testing under 10% ethanol, water, and 20% sucrose conditions. The CSI quantified relative spatial allocation to the Social Zone versus the combined nonsocial zones. Exploratory baseline stratification identified low-CSI (*n* = 7) and high-CSI (*n* = 8) groups, which remained fixed throughout subsequent analyses. Voluntary ethanol consumption and preference were assessed using two-bottle choice with 3% and 6% ethanol. Mice underwent three weeks of chronic intermittent ethanol vapor exposure, followed by post-exposure SACT testing beginning 24 hours after the final vapor exposure. The same animals were followed longitudinally across baseline and post-exposure assessments.

**Results:** At baseline, mice spent more time in the Social Zone than in the Bottle Zone, and CSI differed across bottle conditions. Baseline low-CSI and high-CSI groups showed no significant differences in voluntary ethanol consumption or preference. Following chronic ethanol exposure, whole-cohort ethanol testing decreased Social Zone time and increased Bottle Zone time; whole-cohort water testing showed the opposite pattern, whereas neither measure changed significantly during whole-cohort sucrose testing. CSI-stratified analyses revealed additional heterogeneity in these longitudinal patterns. In high-CSI mice, ethanol testing was associated with decreased Social Zone time, increased Bottle Zone time, and reduced CSI, whereas water testing produced the opposite pattern. Water CSI increased from baseline in both CSI groups, whereas sucrose CSI increased significantly only in high-CSI mice. Across individual animals, greater 6% ethanol preference was associated with a larger baseline-to-post-exposure increase in ethanol-condition Bottle Zone time and a larger decrease in Social Zone time.

**Conclusions:** The SACT detected divergent longitudinal patterns in social-versus-bottle spatial allocation across ethanol, water, and sucrose testing. CSI stratification provided additional insight into behavioral heterogeneity, while individual longitudinal changes in ethanol-condition zone occupancy covaried with separately measured ethanol preference. These findings position the SACT as a complementary framework for quantifying behavioral allocation not captured by ethanol consumption or preference measures alone.

## Introduction

Alcohol use disorder (AUD) is characterized by impaired control over alcohol consumption and may involve diminished engagement with alternative rewarding activities (Sebold et al., 2022). Behavioral-economic models of substance use disorders emphasize both consumption and the allocation of behavior toward drug-related rewards at the expense of available alternatives (Bickel et al., 2014). Preclinical choice procedures underscore the importance of examining these dimensions concurrently. For example, in an alcohol-versus-saccharin procedure, most rats opted for the nondrug reward, whereas a minority persistently chose alcohol and exhibited elevated motivation for it, continuing to seek alcohol despite adverse consequences (Augier et al., 2018). These findings underscore the importance of assessing behavioral allocation—the distribution of behavior among available options—in addition to consumption. A longitudinal approach further extends this framework by characterizing how allocation shifts relative to each animal’s baseline, preserving individual trajectories that may not be evident from group-level analyses.

Social experience serves as a reinforcing stimulus in mice, supporting the inclusion of social stimuli as alternatives to alcohol-associated options (Dölen et al., 2013). However, the relationship between social behavior and alcohol use is complex. In humans, experimental alcohol administration has been shown to enhance positive affect and social bonding during interactions among unfamiliar individuals (Sayette et al., 2012). Social context also influences drinking behavior. Longitudinal social-network analyses have revealed clustering of alcohol consumption among socially connected individuals, although such observational associations do not disentangle interpersonal influence from selection effects and shared exposures (Rosenquist et al., 2010). Experimental studies further indicate that the drinking behavior of others can modify an individual’s consumption, supporting social modeling as a mechanism through which social context affects alcohol use (Collins et al., 1985, Larsen et al., 2012).

Preclinical studies similarly demonstrate that social experience and alcohol-related behavior can influence each other, with effects contingent on experimental conditions. Social isolation has been shown to increase ethanol intake in male mice (Patel et al., 2026), while the impact of chronic ethanol exposure on subsequent social behavior varies across mouse strains. For instance, withdrawal from chronic intermittent ethanol exposure reduced sociability in DBA/2J males but not in C57BL/6J males under the conditions examined by Sidhu et al. (2018). Collectively, these findings highlight the relevance of social stimuli to alcohol-related behavioral allocation and caution against assuming a uniform relationship between ethanol exposure, social approach, and drinking across different experimental contexts.

Established preclinical procedures assess distinct aspects of alcohol-related behavior. Two-bottle choice paradigms measure voluntary ethanol consumption and preference (Xiao et al., 2023), drinking-in-the-dark procedures evaluate binge-like ethanol intake (Rhodes et al., 2005), and operant self-administration procedures quantify responding for ethanol access or delivery (Marchant et al., 2023). However, without the inclusion of competing alternatives, these approaches do not directly assess allocation between alcohol-associated and social options. The presence of alternatives also influences the behavioral consequences of ethanol exposure. For example, in a concurrent operant procedure offering sweetened ethanol and sucrose, chronic ethanol vapor exposure reduced ethanol-directed choice and intake in both male and female rats (Colarusso and Weiner, 2026).Therefore, vapor exposure does not increase ethanol-directed allocation when an alternative reward is available. This context dependence underscores the importance of incorporating alternative options when characterizing alcohol-related behavior, rather than inferring allocation solely from exposure history or consumption.

Direct comparisons of alcohol- and social-directed behavior form the basis for this approach. In operant discrete-choice procedures, rats often selected alcohol over access to a social partner, although allocation varied depending on response requirements and other experimental conditions (Marchant et al., 2023, Augier et al., 2023). In mice, Su et al. (2023) adapted a three-chamber apparatus to assess behavior toward alcohol access and a social stimulus within the same environment, following characterization of alcohol preference using two-bottle choice. These studies establish complementary methodologies for examining alcohol-associated and social behavior within a shared choice context. Building on this work, an important question is how social-versus-bottle spatial allocation shifts relative to each animal’s baseline across an ethanol-exposure period and under different bottle conditions. Repeated spatial assessments enable characterization of these longitudinal patterns at both the cohort and individual-animal level. Individual differences in baseline social approach contribute to heterogeneity within this longitudinal framework. Human studies indicate that sociability-related traits, such as extraversion, are linked to alcohol use, though these associations tend to be modest and vary depending on context. Higher extraversion has been linked to heavy drinking and transitions from moderate to heavy consumption in large adult samples (Hakulinen et al., 2015), as well as to greater alcohol-related mood enhancement in experimental social settings (Fairbairn et al., 2015). In an experience-sampling study of college students, social and enhancement motives statistically mediated associations between extraversion and alcohol outcomes (Hussong, 2003).

A review of drinking motives among adolescents and young adults similarly identified social motives as commonly endorsed reasons for drinking, while enhancement motives were more closely associated with heavy consumption (Kuntsche et al., 2005). These findings justify examining individual behavioral variation but do not imply equivalence between human extraversion and spatial social approach in mice. Consistent with the importance of context, a multivariable behavioral profile including lower sociability predicted greater ethanol preference in adolescent mice, whereas the corresponding adult model did not reliably predict preference (Portugal et al., 2026). Baseline social approach may therefore serve as a useful dimension for exploring heterogeneity, without suggesting that sociability is a universal or deterministic predictor of alcohol-related outcomes.

Three-chamber procedures evaluate social approach by comparing exploration of social and nonsocial locations (Nadler et al., 2004). However, chamber occupancy should be distinguished from direct social investigation or sustained affiliative interaction (Sofyana et al., 2026). In the present assay, an adapted Chamber Sociability Index (CSI) summarizes time spent in the Social Zone relative to the combined Neutral and Bottle Zones. This index reflects social-versus-combined-nonsocial spatial allocation rather than directly measuring ethanol preference. Similarly, Bottle Zone occupancy alone does not quantify fluid consumption or establish reinforcement. Within this measurement framework, baseline CSI can be used descriptively to investigate whether animals with different initial allocation patterns display heterogeneous longitudinal trajectories. These classifications are exploratory behavioral strata rather than distinct biological subtypes or validated markers of alcohol vulnerability.

In this study, a modified three-chamber assay, termed the Social Alcohol Choice Test (SACT), was developed to characterize longitudinal changes in spatial allocation between a caged, non-cagemate male conspecific and a bottle-associated option, separated by a neutral central zone. The same 15 male C57BL/6J mice were tested at baseline and after 3 weeks of chronic intermittent ethanol (CIE) vapor exposure, with post-exposure testing commencing 24 hours after the final exposure. Separate sessions with 10% ethanol, water, or 20% sucrose enabled comparisons across bottle conditions. The primary objective was to characterize within-animal changes in Social and Bottle Zone occupancy and to examine how longitudinal patterns varied across these conditions. Additionally, the study assessed whether individual baseline-to-post-exposure changes during ethanol testing were associated with 6% ethanol preference measured separately by two-bottle choice during a vapor-free interval. For exploratory analysis, mice were stratified by their mean baseline CSI across the three bottle conditions, with low-CSI and high-CSI assignments maintained throughout follow-up. Additional measures included systemic ethanol exposure, open-field center exploration, and locomotor activity. This longitudinal framework was designed to complement consumption-based measures by describing changes in spatial allocation within the same animals and exploring their relationship to separately measured ethanol preference.

## Materials and Methods

### Animals

Six-week-old male C57BL/6J mice (*n* = 15) were purchased from The Jackson Laboratory (Bar Harbor, ME) and group-housed five per cage in standard Plexiglas cages at Mayo Clinic (Rochester, MN). The colony room was maintained at 24 ± 1°C and 60% ± 2% relative humidity under a 12-hour light/dark cycle, with lights on at 06:00 and lights off at 18:00. Mice had *ad libitum* access to food and water throughout the study. Following arrival, mice were allowed to acclimate to the animal facility for 1 week and were subsequently handled for 1 week before behavioral testing. Behavioral experiments were conducted during the light phase.

Separate male mice served as non-cagemate social stimulus animals during the Social Alcohol Choice Test (SACT). Stimulus mice were housed separately from the experimental mice and were not included in behavioral or statistical analyses. All experimental procedures were approved by the Mayo Clinic Institutional Animal Care and Use Committee and were conducted in accordance with National Institutes of Health guidelines for the care and use of laboratory animals.

### Experimental design

The study was designed primarily as a longitudinal within-subject experiment. The same 15 experimental mice underwent baseline SACT testing, chronic intermittent ethanol (CIE) vapor exposure, and post-exposure SACT testing. Each animal was assessed under 10% ethanol, water, and 20% sucrose bottle conditions, allowing longitudinal changes in behavioral allocation to be evaluated relative to each mouse’s own baseline. Animal identity was retained across repeated bottle conditions and baseline and post-exposure assessments.

Baseline variation in Chamber Sociability Index (CSI) was subsequently examined as an exploratory source of behavioral heterogeneity. Exploratory CSI stratification was therefore secondary to the primary longitudinal design and did not represent an experimentally assigned treatment condition. Low-CSI and high-CSI classifications were fixed relative to the baseline and retained in subsequent analyses.

### Chronic intermittent ethanol vapor exposure

All experimental mice underwent CIE vapor exposure using a modified vapor-chamber procedure based on established CIE methods and protocols previously used in our laboratory (Hong et al., 2019, Shang et al., 2020, Peyton et al., 2025, Kang et al., 2025). Mice remained in their home cages within custom-built Plexiglas vapor chambers designed and constructed by Mayo Clinic engineering staff and were exposed to ethanol vapor for 16 hours per day, followed by an 8-hour period without vapor. Vapor exposure occurred on 4 consecutive days followed by 3 days without vapor, and this weekly exposure schedule was repeated for 3 weeks.

Ethanol vapor was generated by passing air through a flask containing 95% ethanol at a flow rate of 200–250 mL/min and mixing the ethanol-containing air with room air before delivery to the vapor chamber. Ethanol exposure was progressively increased during the first 2 exposure days of each week and maintained at a consistent level during exposure days 3 and 4. Chamber conditions were adjusted to achieve a target blood alcohol concentration (BAC) of 250–300 mg/dL. In contrast to some previously published CIE protocols, mice did not receive ethanol loading doses or pyrazole before vapor exposure.

During the third week of CIE exposure, approximately 50 µL of blood was collected from each experimental mouse in the procedure room adjacent to the vapor chamber room approximately 5–15 minutes after removal from the ethanol vapor chamber following completion of the daily vapor exposure. BAC was measured using an Analox GL5 multimetabolite analyzer (Analox Instruments, Stourbridge, United Kingdom). These measurements were used to characterize systemic ethanol exposure during CIE.

### Social Alcohol Choice Test

Behavioral allocation between a social stimulus and a bottle-associated option was assessed using a modified three-chamber Social Alcohol Choice Test (SACT). Testing was performed in a Panlab LE894 three-compartment social interaction apparatus (Panlab, Barcelona, Spain; serial no. 3013620) measuring 60 × 42 × 22 cm, with each compartment measuring 20 × 42 × 22 cm. One lateral chamber, designated the Social Zone, contained a non-cagemate male stimulus mouse confined within a wire enclosure. The opposite lateral chamber, designated the Bottle Zone, contained a bottle equipped with a sipper tube connected to a small drinking trough. The center chamber served as the Neutral Zone and contained no social or bottle stimulus.

The bottle contained 10% ethanol, water, or 20% sucrose depending on the test condition. The bottle assembly consisted of a 50-mL conical tube fitted with a straight 1.5-inch stainless-steel ball-bearing sipper tube and a black rubber No. 8.5 bottle stopper. The sipper tube delivered fluid to a small drinking trough accessible to the experimental mouse within the Bottle Zone. Although mice could access the solution during testing, the primary SACT outcomes quantified spatial allocation within the apparatus rather than fluid consumption.

Testing was conducted concurrently in two behavioral testing rooms by two experimenters. Seven mice were consistently tested in Room A by Experimenter A and eight mice in Room B by Experimenter B. Each experimental mouse retained the same room and experimenter assignment during baseline and post-exposure testing to maintain consistency across longitudinal assessments. [INSERT approximate illumination and SACT testing time relative to the light cycle.]

At the beginning of each trial, the experimental mouse was placed in the Neutral Zone, and the 15-minute test began immediately; no separate habituation period was provided. Mice were allowed to freely explore all three zones for the duration of the trial and were removed from the apparatus after 15 minutes. Stimulus mice were rotated after every two experimental mice. Between trials, the apparatus and associated testing components were cleaned with 70% ethanol and allowed to dry before the next mouse was tested.

Baseline SACT testing was performed before CIE exposure over 6 consecutive test days. Mice were tested with 10% ethanol on days 1 and 2, water on days 3 and 4, and 20% sucrose on days 5 and 6. For each bottle condition, the positions of the social stimulus and bottle were reversed on the second test day to assess and control for potential side preference.

Following 3 weeks of CIE exposure, post-exposure SACT testing began 24 hours after the final ethanol vapor exposure. Post-exposure testing was conducted on 3 consecutive days in the fixed order of 10% ethanol, water, and 20% sucrose, with one 15-minute trial performed for each condition. No evidence of systematic side preference was detected during baseline testing when the Social and Bottle Zone positions were reversed. Accordingly, post-exposure Social and Bottle Zone positions were assigned by the experimenter and were not further counterbalanced across test days.

Behavior was recorded and analyzed using EthoVision XT 14 (Noldus Information Technology, Wageningen, the Netherlands). The Social, Neutral, and Bottle Zones were defined according to the physical boundaries of the three apparatus compartments. Zone occupancy was determined by tracking the experimental mouse’s nose point within the predefined zone boundaries. Total time spent in each zone, distance traveled, and transitions between the Social and Bottle Zones were quantified.

Relative spatial allocation toward the social stimulus was additionally quantified using a Chamber Sociability Index (CSI) adapted from Sofyana et al. (2026). In the current SACT paradigm, nonsocial time was defined as the sum of time spent in the Neutral and Bottle Zones. CSI was calculated as:

CSI = [Social Zone time − (Neutral Zone time + Bottle Zone time)] / [Social Zone time + Neutral Zone time + Bottle Zone time]

CSI values range from −1 to +1, with positive values indicating greater relative allocation to the Social Zone and negative values indicating greater relative allocation to the combined Neutral and Bottle Zones.

For exploratory analyses of baseline behavioral heterogeneity, a mean baseline CSI was calculated for each mouse by averaging its CSI values across the 10% ethanol, water, and 20% sucrose baseline conditions. Mice were then assigned to fixed low-CSI (*n* = 7) and high-CSI (*n* = 8) groups according to the sign of this mean baseline CSI. Mice with a positive mean baseline CSI (CSI > 0) were assigned to the high-CSI group, whereas mice with a negative mean baseline CSI (CSI < 0) were assigned to the low-CSI group. These classifications were retained for all subsequent analyses, allowing longitudinal behavior to be evaluated within the same baseline-defined CSI strata rather than redefining groups after CIE exposure.

### Open-field test

Open-field testing was conducted in an ENV-510 open-field test environment (Med Associates, St. Albans, VT, USA) consisting of a 27 × 27 × 20.3 cm Plexiglas arena equipped with infrared photobeams. Locomotor activity was recorded using Activity Monitor software (Med Associates).

Two open-field test (OFT) sessions were included in the analyses. The first was conducted at baseline before CIE exposure and the second at the end of Week 2 of CIE exposure. Both sessions were performed between approximately 1:00 and 2:00 PM. An additional OFT session was attempted during Week 3; however, the data were excluded due to technical issues during testing.

Time spent in the center zone was quantified in seconds to measure exploration of the center. The center zone was defined as the central 13 × 13 cm region of the arena, consistent with previously used open-field definitions in our laboratory (Peyton et al., 2025, Ruby et al., 2011). Locomotor activity was quantified as distance traveled in centimeters. Although mice were allowed to explore the arena for 30 minutes, analyses reported in Figure 3C–D were restricted to the first 10 minutes of each session. Center-zone time and distance traveled during this interval were quantified using Activity Monitor software. Between animals, the open-field arena was cleaned with 70% ethanol and allowed to dry before the next test.

### Two-bottle choice ethanol drinking

Voluntary ethanol consumption and preference were assessed using a two-bottle choice (TBC) procedure during designated 72-hour vapor-free periods between CIE exposure cycles, following established procedures used in our laboratory (Hong et al., 2019, Peyton et al., 2025, Peyton et al., 2022). At the end of Week 1 of CIE exposure, mice were individually housed and given continuous access to 3% ethanol and water for 72 hours. At the end of Week 2, the procedure was repeated using 6% ethanol and water. Food remained available *ad libitum* throughout both assessments.

Bottle positions were reversed daily to minimize side preference. Mice and bottles were weighed at the beginning and end of each 72-hour test period. Identical bottles were placed on two empty cages during testing to estimate fluid loss attributable to evaporation or leakage. Ethanol solution consumption was corrected by subtracting the mean fluid loss measured from the control bottles.

Ethanol intake was expressed as grams of ethanol consumed per kilogram of body weight (g/kg) and calculated as:

Ethanol intake (g/kg) = [corrected ethanol solution consumed (mL) × ethanol concentration (v/v) × 0.789 g/mL] / body weight (kg) where 0.789 g/mL represents the density of ethanol.

### Statistical analysis

Statistical analyses were performed using GraphPad Prism 11 for Windows (GraphPad Software, Boston, MA, USA). Statistical tests were two-tailed unless otherwise specified, and α was set at 0.05. Data are presented as mean ± SEM unless otherwise indicated.

The primary SACT analyses used a longitudinal repeated-measures framework in which animal identity was retained across measurements. The same 15 mice contributed measurements across bottle conditions and baseline and post-exposure sessions. Accordingly, bottle condition (10% ethanol, water, and 20% sucrose), testing session (baseline and post-exposure), and zone, when included in the same model, were treated as within-subject factors. Comparisons between low-CSI and high-CSI mice represented the only between-subject comparisons in the SACT analyses. Within the low-CSI and high-CSI groups, baseline-to-post-exposure and across-condition comparisons remained paired because the same animals were repeatedly measured.

Baseline CSI stratification was treated as an exploratory analysis of behavioral heterogeneity rather than an experimentally assigned treatment factor. Low-CSI and high-CSI classifications were defined from each animal’s mean CSI across the three baseline bottle conditions and remained fixed throughout all subsequent analyses.

Depending on the experimental design, comparisons were performed using paired or unpaired *t* tests, one-way repeated-measures ANOVA, two-way repeated-measures ANOVA, or mixed-design ANOVA. Geisser–Greenhouse correction was applied when sphericity was not assumed.

Multiple comparisons were performed using Tukey, Šídák, or Fisher’s least significant difference (LSD) procedures, as appropriate for each analysis. Welch’s correction was applied to unpaired t-tests when unequal variances were present.

Simple linear regression was used for the BAC calibration curve and to evaluate associations between 6% ethanol preference and individual baseline-to-post-exposure changes in ethanol-condition SACT zone occupancy. These regression analyses assessed covariation between continuous measures and were not interpreted as establishing causality. The specific statistical tests, degrees of freedom, multiple-comparison procedures, exact *P* values, confidence intervals, and effect sizes, when applicable, are reported with the corresponding Results and figure analyses.

## Results

### Baseline SACT behavior favored the Social Zone, with CSI differing across bottle conditions

The Social–Alcohol Choice Test (SACT) was used to assess competition between social interaction and access to a bottle containing 10% ethanol, water, or 20% sucrose (Figure 1A). The overall experimental sequence, including baseline SACT testing, chronic intermittent ethanol (CIE) exposure, open field testing, and post-CIE SACT assessment, is shown in Figure 1B.

**Figure 1.**
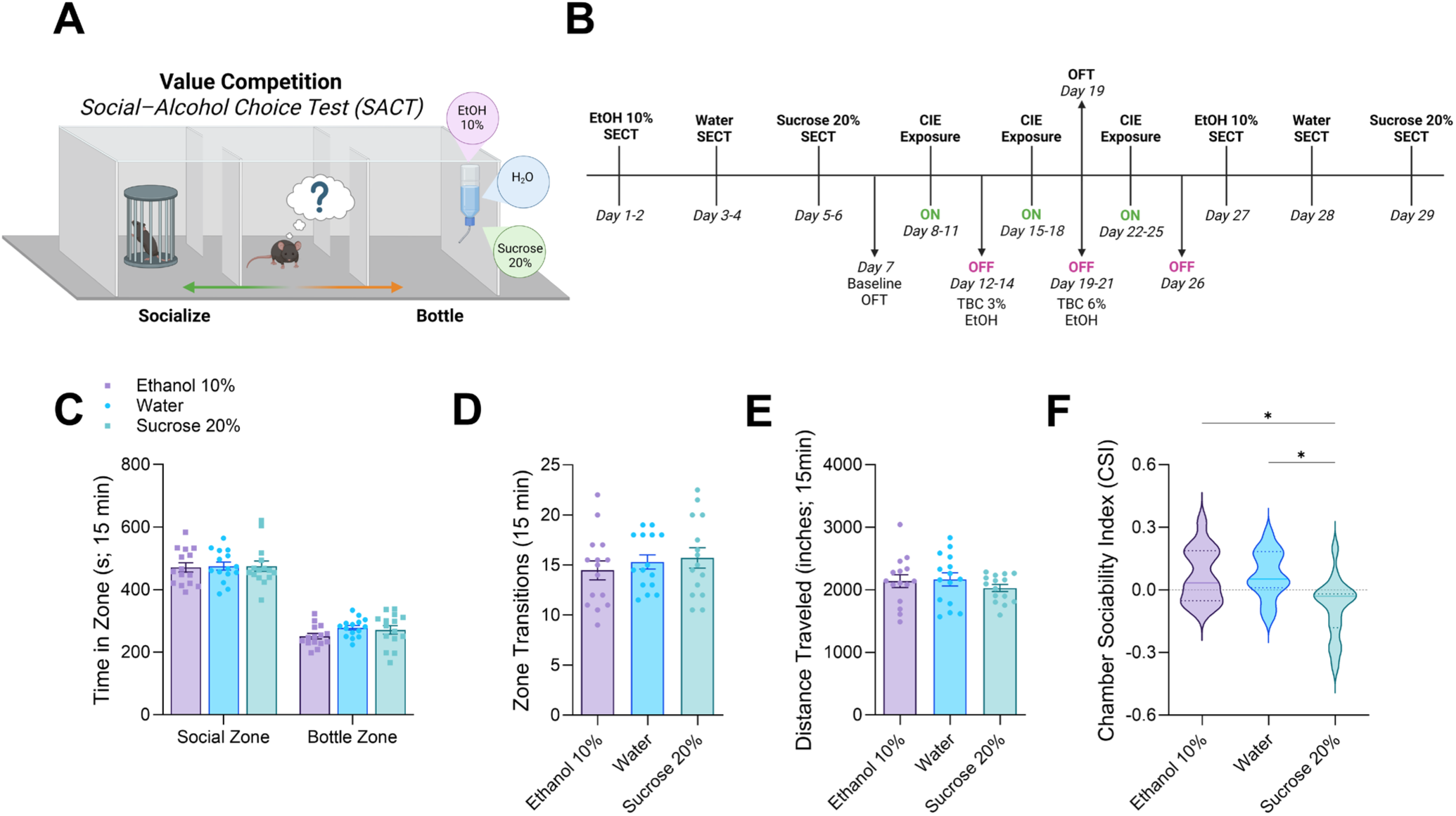
Baseline Social–Alcohol Choice Test behavior and experimental timeline. (A) Schematic of the Social–Alcohol Choice Test (SACT), in which mice choose between proximity to a non-cagemate male social stimulus in the Social Zone and a bottle containing 10% ethanol, water, or 20% sucrose in the Bottle Zone. (B) Experimental timeline showing baseline SACT testing with 10% ethanol (Days 1–2), water (Days 3–4), and 20% sucrose (Days 5–6), followed by baseline open field testing (OFT; Day 7), chronic intermittent ethanol (CIE) exposure, two-bottle choice (TBC) testing during CIE OFF periods, and post-CIE SACT testing with 10% ethanol, water, and 20% sucrose on Days 27–29, respectively. (C) Time spent in the Social Zone and Bottle Zone during baseline SACT testing across bottle conditions. (D) Number of zone transitions during baseline SACT testing. (E) Distance traveled during the 15-minute baseline SACT session. (F) Chamber Sociability Index (CSI) across the three baseline bottle conditions. The same 15 mice were tested under all three bottle conditions. Individual animals are shown in C–E; bars represent mean ± SEM. Violin plots in F show the distribution of individual CSI values with median and quartiles. Baseline zone occupancy was analyzed by two-way repeated-measures ANOVA with zone and bottle condition as within-subject factors, followed by Tukey-adjusted paired comparisons. CSI, distance traveled, and zone transitions were analyzed using one-way repeated-measures ANOVA with a Greenhouse–Geisser correction when sphericity was not assumed, and Tukey-adjusted paired comparisons. *P* < 0.05. EtOH, ethanol; H₂O, water; SACT, Social–Alcohol Choice Test; CIE, chronic intermittent ethanol; TBC, two-bottle choice; OFT, open field test; CSI, Chamber Sociability Index

Baseline SACT behavior was examined in the same 15 mice under conditions 10% ethanol, water, and 20% sucrose. Because each animal was tested under all three bottle conditions, bottle condition was treated as a within-subject factor. Social Zone and Bottle Zone measurements were also repeated within each animal. A two-way repeated-measures ANOVA with zone and bottle condition as within-subject factors showed greater time in the Social Zone than in the Bottle Zone when averaged across conditions (*F*(1.000, 14.00) = 150.2, *P* < 0.0001; mean difference, 207.0 s; 95% CI, 170.8 to 243.2). Neither the main effect of bottle condition (*F*(1.597, 22.36) = 2.954, *P* = 0.0825) nor the zone × condition interaction (*F*(1.520, 21.28) = 0.3684, *P* = 0.6389) was significant. Tukey-adjusted paired comparisons within zones showed lower Bottle Zone time during the 10% ethanol condition than during the water condition (mean difference, −27.19 s; 95% CI, −45.74 to −8.645; adjusted *P* = 0.0048), whereas the remaining Bottle Zone comparisons were not significant. Social Zone time did not differ between ethanol and water, ethanol and sucrose, or water and sucrose conditions (adjusted *P* = 0.9627, 0.9840, and 0.9996, respectively; **Figure 1C**).

Chamber Sociability Index (CSI) was compared across the three bottle conditions using repeated measurements from the same animals and differed significantly across conditions (Geisser– Greenhouse-corrected one-way repeated-measures ANOVA, *F*(1.549, 21.69) = 6.489, *P* = 0.0098). Tukey-adjusted paired comparisons showed lower CSI during sucrose testing than during ethanol or water testing (adjusted *P* = 0.0250 and 0.0485, respectively), whereas ethanol and water did not differ (adjusted *P* = 0.9975; **Figure 1F**).

Baseline distance traveled did not differ significantly across ethanol, water, and sucrose conditions (Geisser–Greenhouse-corrected one-way repeated-measures ANOVA, *F*(1.327, 18.57) = 1.341, *P* = 0.2736; **Figure 1E**). Consistent with the omnibus analysis, Tukey-adjusted paired comparisons detected no differences between ethanol and water (adjusted *P* = 0.8522), ethanol and sucrose (adjusted *P* = 0.5287), or water and sucrose (adjusted *P* = 0.4218). The number of zone transitions likewise did not differ significantly across baseline bottle conditions (Geisser–Greenhouse-corrected one-way repeated-measures ANOVA, *F*(1.993, 27.90) = 0.9040, *P* = 0.4162; Figure 1D).

### Exploratory baseline CSI stratification identified heterogeneity in ethanol-condition zone occupancy

Because baseline CSI varied among individual mice, we next explored whether this variation characterized heterogeneity in SACT behavior. A mean baseline CSI was calculated for each mouse by averaging its CSI values across the 10% ethanol, water, and 20% sucrose baseline conditions. Based on the sign of this mean baseline CSI, mice were assigned to fixed low-CSI (*n* = 7) and high-CSI (*n* = 8) strata that were retained for subsequent exploratory analyses. Bottle condition remained a within-subject factor, whereas CSI stratum represented a between-subject factor when the two strata were compared.

Across bottle conditions, CSI showed a significant main effect of condition (*F*(1.420, 18.46) = 6.703, *P* = 0.0114) and, as expected because the strata were defined from the same baseline CSI measurements, a significant main effect of CSI stratum (*F*(1, 13) = 14.16, *P* = 0.0024). The condition × CSI-stratum interaction was not significant (*F*(1.420, 18.46) = 0.9598, *P* = 0.3722). Tukey-adjusted comparisons showed lower CSI in low-CSI than high-CSI mice during ethanol testing (adjusted *P* = 0.0033), whereas the strata did not differ significantly during water or sucrose testing (adjusted *P* = 0.3043 and 0.5091, respectively). Within low-CSI mice, paired comparisons across bottle conditions showed lower CSI during ethanol than water testing (adjusted *P* = 0.0432), whereas ethanol versus sucrose and water versus sucrose comparisons were not significant (adjusted *P* = 0.3380 and 0.1541, respectively; Figure 2A).

**Figure 2.**
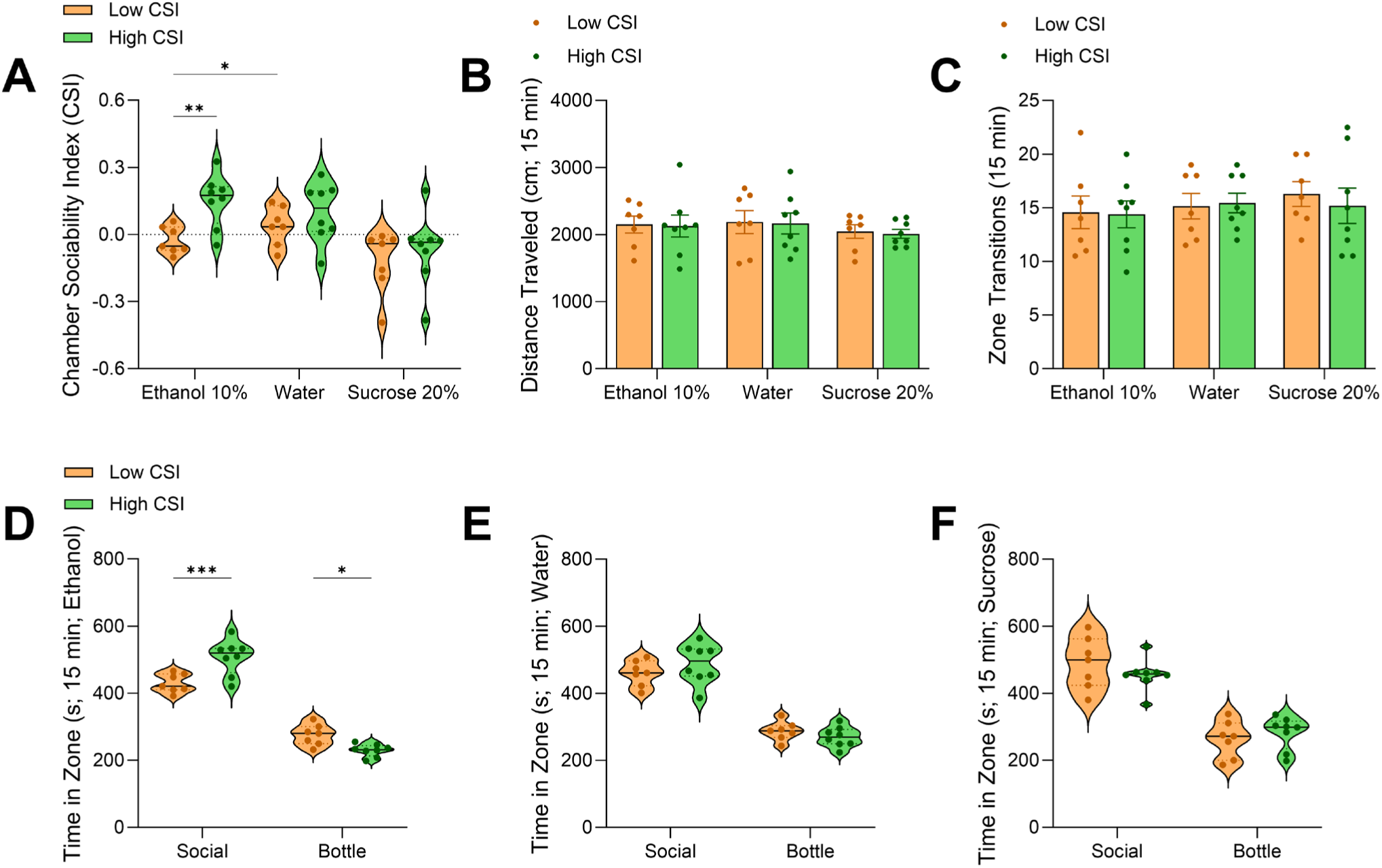
Baseline SACT behavior in exploratory low- and high-CSI strata. Mice were stratified according to the sign of their mean baseline Chamber Sociability Index (CSI), calculated across the 10% ethanol, water, and 20% sucrose conditions of the Social–Alcohol Choice Test (SACT), yielding fixed low-CSI (*n* = 7) and high-CSI (*n* = 8) strata. (A) CSI across the three baseline bottle conditions in low- and high-CSI mice. (B) Distance traveled during the 15-minute SACT session across bottle conditions. (C) Number of zone transitions during the 15-minute SACT session. (D–F) Time spent in the Social Zone and Bottle Zone by low- and high-CSI mice during (D) 10% ethanol, (E) water, and (F) 20% sucrose conditions. Individual animals are shown. Bars in B and C represent mean ± SEM; violin plots in A and D–F show the distributions of individual values with median and quartiles. CSI, distance traveled, and zone transitions were analyzed by two-way repeated-measures ANOVA with bottle condition as a within-subject factor and baseline CSI stratum as a between-subject factor. Zone occupancy was analyzed separately for each bottle condition by two-way repeated-measures ANOVA with zone as a within-subject factor and CSI stratum as a between-subject factor. For the 10% ethanol condition, the significant zone × CSI-stratum interaction was followed by uncorrected Fisher’s least significant difference comparisons of low-versus high-CSI mice within each zone. For the water and 20% sucrose conditions, the zone × CSI-stratum interactions were not significant; Šídák-adjusted comparisons of low-versus high-CSI mice within each zone were also not significant. *P* < 0.05, **P** < 0.01, ***P*** < 0.001. SACT, Social–Alcohol Choice Test; CSI, Chamber Sociability Index.

**Figure 3.**
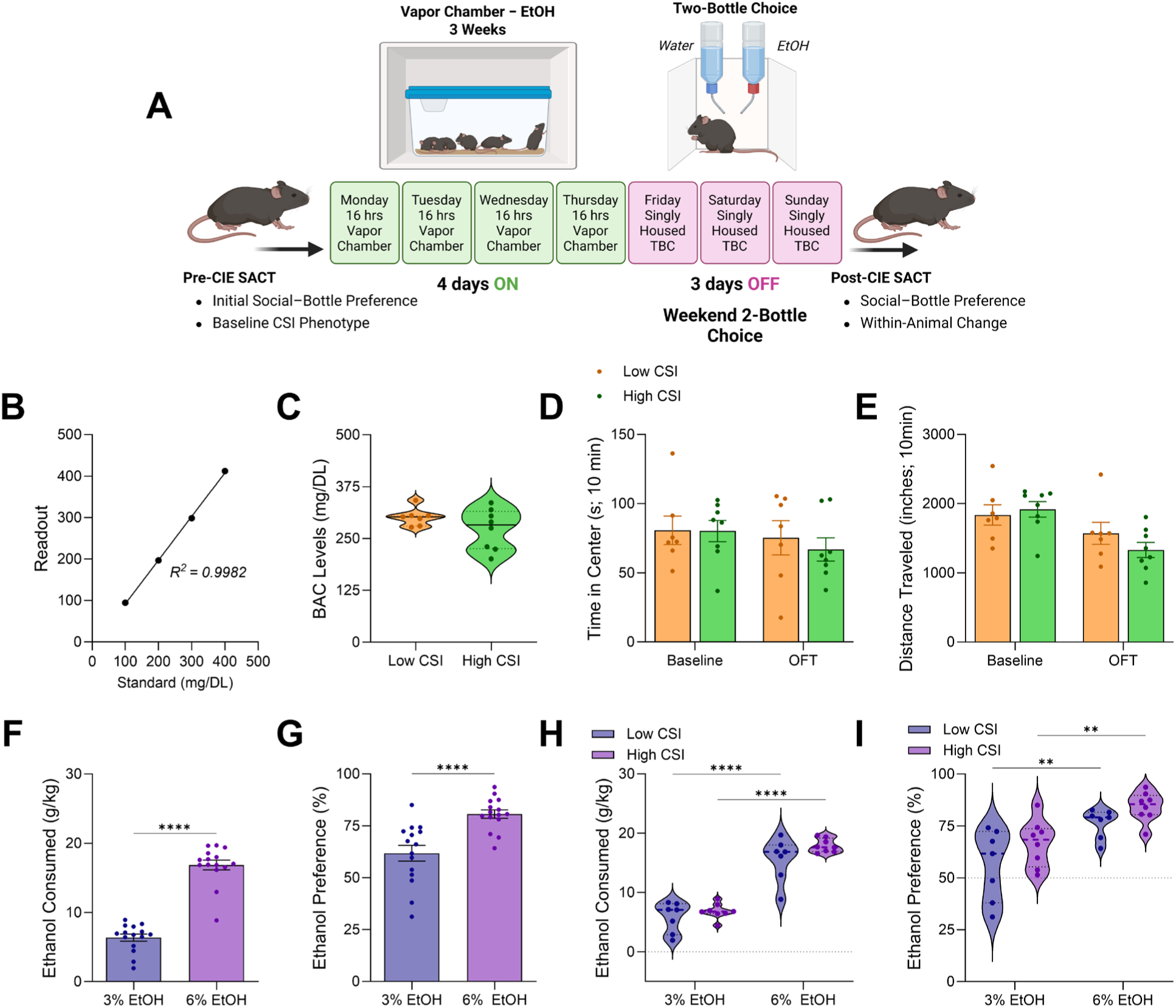
Chronic intermittent ethanol exposure and voluntary ethanol drinking in baseline-defined CSI strata. (A) Schematic of the chronic intermittent ethanol (CIE) paradigm consisting of 4 consecutive days of 16-hour ethanol vapor exposure followed by 3 days OFF, during which two-bottle choice (TBC) testing was performed. Low-CSI (*n* = 7) and high-CSI (*n* = 8) strata were defined from pre-CIE Social–Alcohol Choice Test (SACT) behavior and remained fixed throughout subsequent analyses. (B) Blood alcohol concentration (BAC) assay calibration curve. (C) BAC levels in low- and high-CSI mice. (D) Open-field center time and (E) distance traveled at baseline and during CIE exposure. (F) Ethanol consumption and (G) ethanol preference during 3% and 6% ethanol TBC testing across all mice (*n* = 15). (H) Ethanol consumption and (I) ethanol preference during 3% and 6% ethanol access in low- and high-CSI mice. Individual animals are shown; bars represent mean ± SEM, and violin plots show individual distributions with median and quartiles. B was analyzed by simple linear regression; C by Welch-corrected unpaired *t* test; D and E by two-way repeated-measures ANOVA; F and G by paired *t* tests; and H and I by two-way repeated-measures ANOVA with Fisher’s LSD (H) or Šídák-adjusted (I) comparisons. **P** < 0.01; **P** < 0.0001. CIE, chronic intermittent ethanol; SACT, Social–Alcohol Choice Test; TBC, two-bottle choice; CSI, Chamber Sociability Index; BAC, blood alcohol concentration.

Baseline distance traveled was likewise assessed repeatedly in the same animals across the three bottle conditions. There was no significant main effect of CSI stratum (*F*(1, 13) = 0.02955, *P* = 0.8662), main effect of bottle condition (*F*(1.376, 17.88) = 1.288, *P* = 0.2869), or condition × CSI-stratum interaction (*F*(1.376, 17.88) = 0.003198, *P* = 0.9838; Figure 2B). Baseline transition counts similarly showed no significant condition × CSI-stratum interaction (*F*(1.991, 25.88) = 0.2688, *P* = 0.7654), main effect of condition (*F*(1.991, 25.88) = 0.8843, *P* = 0.4247), or main effect of CSI stratum (*F*(1, 13) = 0.05035, *P* = 0.8260; Figure 2C).

Zone occupancy was analyzed using repeated Social Zone and Bottle Zone measurements within each mouse, with CSI stratum as the between-subject factor. During the ethanol condition, there was a significant zone × CSI-stratum interaction (*F*(1, 13) = 15.68, *P* = 0.0016). Uncorrected

Fisher’s least significant difference comparisons showed that low-CSI mice spent less time in the Social Zone than high-CSI mice (estimated mean difference, low minus high, −77.96 s; 95% CI, −115.1 to −40.80; *P* = 0.0002) and more time in the Bottle Zone (estimated mean difference, 46.04 s; 95% CI, 8.888 to 83.20; *P* = 0.0171; Figure 2D). In contrast, the zone × CSI-stratum interaction was not significant during water (*F*(1, 13) = 1.539, *P* = 0.2366) or sucrose testing (*F*(1, 13) = 1.014, *P* = 0.3323). Šídák-adjusted comparisons detected no between-stratum differences in Social or Bottle Zone time during water testing (adjusted *P* = 0.3499 and 0.6617, respectively) or sucrose testing (adjusted *P* = 0.4332 and 0.7651, respectively; Figure 2E,F). Thus, the exploratory baseline CSI stratification identified differences in ethanol-condition spatial allocation that were not accompanied by detectable differences in overall locomotor distance or transition counts.

Supplementary analyses incorporating all three chamber zones further characterized baseline spatial allocation. Within each CSI stratum, both bottle condition and zone were repeated within the same animals. In low-CSI mice, a Geisser–Greenhouse-corrected two-way repeated-measures ANOVA identified a significant zone × bottle-condition interaction (*F*(1.626, 9.754) = 5.221, *P* = 0.0336). Tukey-adjusted paired comparisons showed less Social Zone time during ethanol than water testing (adjusted *P* = 0.0236) and greater Neutral Zone time during ethanol than water or sucrose testing (adjusted *P* = 0.0151 and 0.0015, respectively). Bottle Zone time did not differ significantly across conditions (Figure S1A).

In high-CSI mice, there was a significant main effect of zone (*F*(1.332, 9.322) = 144.1, *P* < 0.0001), whereas the zone × bottle-condition interaction did not reach significance (*F*(1.823, 12.76) = 3.312, *P* = 0.0728). Tukey-adjusted paired comparisons showed greater Social Zone time during ethanol than sucrose testing (adjusted *P* = 0.0335) and lower Bottle Zone time during ethanol than water or sucrose testing (adjusted *P* = 0.0174 and 0.0392, respectively; Figure S1B).

Because the omnibus interaction was not significant, these simple comparisons were not interpreted as demonstrating condition-dependent differences in zone allocation.

### Ethanol exposure, locomotor behavior, and voluntary ethanol drinking were characterized during CIE

The CIE paradigm consisted of four consecutive days of 16-hour ethanol vapor exposure followed by three OFF days, during which two-bottle choice (TBC) testing was performed (Figure 3A).

The BAC assay calibration curve showed a strong linear relationship between standard concentration and assay readout, with a slope of 1.052 (95% CI, 1.023 to 1.081), *R*² = 0.9981, and *F*(1, 12) = 6432, *P* < 0.0001 (**Figure 3B**). Exploratory comparison of systemic ethanol exposure between the baseline-defined CSI strata showed no significant difference in BAC between low-CSI and high-CSI mice (Welch-corrected unpaired *t* test, *t*(9.837) = 1.514, *P* = 0.1615; η² = 0.1890; **Figure 3C**).

Open-field behavior during the first 10 minutes of each session was compared longitudinally between baseline and the end of Week 2 of CIE exposure. The same animals contributed measurements at both sessions; therefore, session was treated as a within-subject factor and baseline CSI stratum as a between-subject factor in the exploratory stratified analysis. Center time showed no significant main effect of session (*F*(1, 13) = 1.193, *P* = 0.2946), CSI stratum (*F*(1, 13) = 0.1787, *P* = 0.6794), or session × CSI-stratum interaction (*F*(1, 13) = 0.2145, *P* = 0.6509; **Figure 3D**). In contrast, distance traveled during the first 10 minutes was lower at the Week 2 assessment than at baseline (main effect of session, *F*(1, 13) = 15.99, *P* = 0.0015), with no significant main effect of CSI stratum (*F*(1, 13) = 0.2728, *P* = 0.6102) or session × CSI-stratum interaction (*F*(1, 13) = 2.280, *P* = 0.1549; **Figure 3E**).

The same 15 mice were tested during both the 3% and 6% ethanol two-bottle choice conditions. Across the cohort, ethanol consumption was higher during access to 6% ethanol than 3% ethanol (paired *t*(14) = 10.79, *P* < 0.0001; partial η² = 0.8926), with a mean within-animal difference of 10.48 g/kg (95% CI, 8.393 to 12.56; **Figure 3F**). In the exploratory CSI-stratified analysis, ethanol concentration was treated as a within-subject factor and baseline CSI stratum as a between-subject factor. There was a significant main effect of ethanol concentration (*F*(1, 13) = 394.6, *P* < 0.0001), but no main effect of CSI stratum (*F*(1, 13) = 2.379, *P* = 0.1470) or concentration × CSI-stratum interaction (*F*(1, 13) = 1.391, *P* = 0.2593). Uncorrected Fisher’s least significant difference comparisons showed greater consumption at 6% than at 3% ethanol within both low-CSI and high-CSI mice (both *P* < 0.0001; **Figure 3H**).

Ethanol preference was likewise higher during access to 6% ethanol than 3% ethanol (paired *t*(14) = 5.915, *P* < 0.0001; partial η² = 0.7142), with a mean within-animal increase of 18.85 percentage points (95% CI, 12.01 to 25.68; Figure 3G). In the exploratory CSI-stratified repeated-measures analysis, there was a significant main effect of ethanol concentration (*F*(1, 13) = 33.00, *P* < 0.0001), with no significant main effect of CSI stratum (*F*(1, 13) = 3.651, *P* = 0.0783) or concentration × CSI-stratum interaction (*F*(1, 13) = 0.1463, *P* = 0.7082). Šídák-adjusted paired comparisons showed higher ethanol preference at 6% than at 3% within both low-CSI and high-CSI mice (adjusted *P* = 0.0021 and 0.0035, respectively; **Figure 3I**).

### SACT zone occupancy and CSI showed condition-specific longitudinal changes following chronic ethanol exposure

The primary longitudinal SACT analyses followed the same 15 mice from baseline through post-exposure testing following 3 weeks of CIE. At both sessions, all mice were tested under 10% ethanol, water, and 20% sucrose conditions; therefore, bottle condition and session were within-subject factors. Zone measurements were likewise repeated within each animal. Exploratory low-CSI (*n* = 7) and high-CSI (*n* = 8) classifications were defined from each mouse’s mean CSI across the three baseline conditions and remained fixed throughout subsequent analyses. The CSI stratum was represented as a between-subjects factor only when the two strata were directly compared.

During the 10% ethanol condition, the whole-cohort zone × session interaction was significant (*F*(1, 14) = 13.63, *P* = 0.0024). Uncorrected Fisher’s least significant difference comparisons showed that Social Zone time decreased from 471.3 s at baseline to 398.4 s post-exposure (*P* = 0.0203), whereas Bottle Zone time increased from 250.9 to 323.4 s (*P* = 0.0208; Figure 4A). When the exploratory baseline CSI strata were examined separately using paired baseline-to-post-exposure comparisons, low-CSI mice showed no significant change in Social Zone time (adjusted *P* = 0.9674) or Bottle Zone time (adjusted *P* = 0.3966). High-CSI mice showed decreased Social Zone time from 507.6 to 380.1 s (adjusted *P* = 0.0080) and increased Bottle Zone time from 229.4 to 339.8 s (adjusted *P* = 0.0047; Figure 4B).

**Figure 4.**
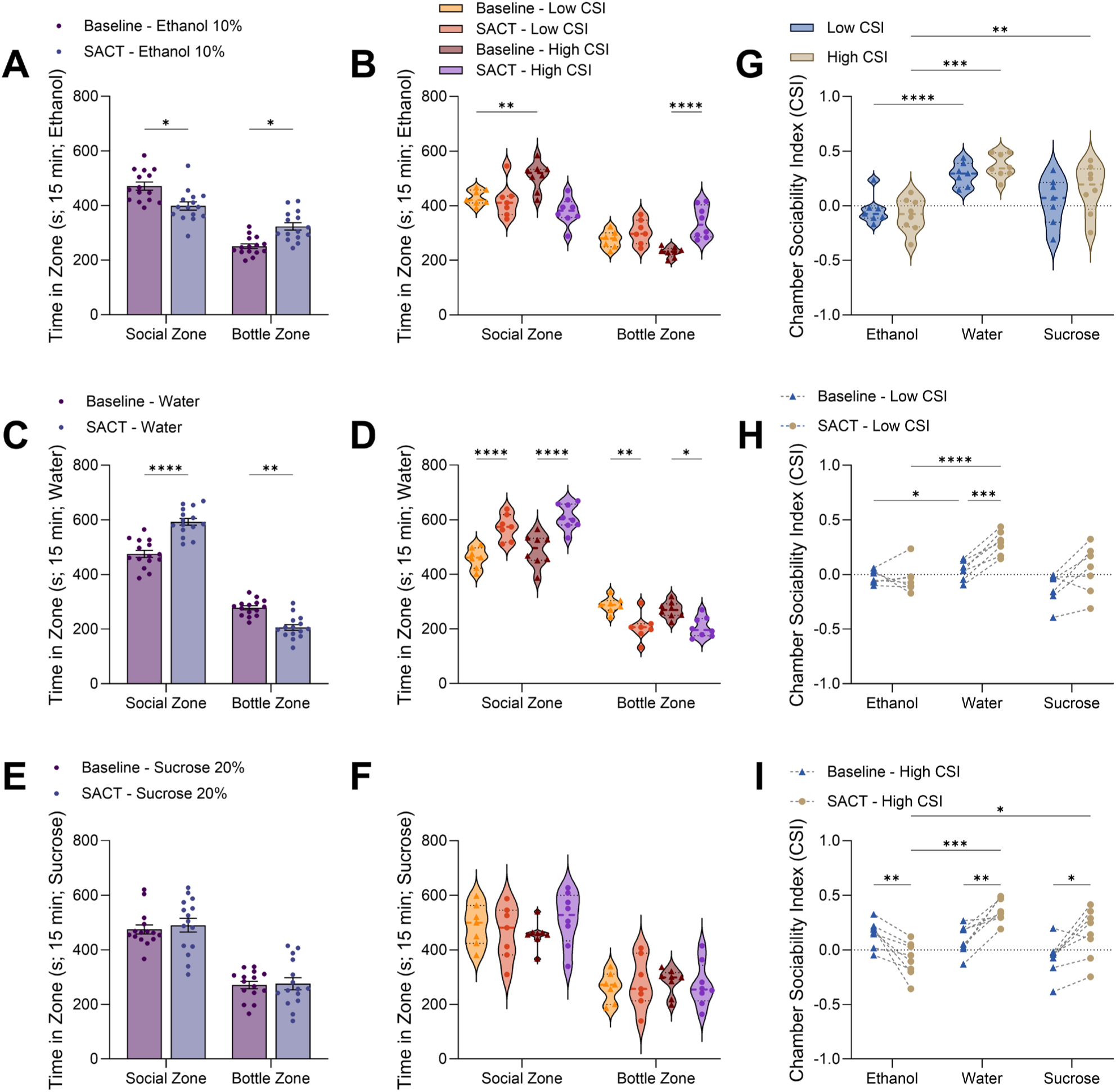
Condition-specific changes in SACT behavior following chronic ethanol exposure. (A, C, E) Social Zone and Bottle Zone time at baseline and post-CIE Social–Alcohol Choice Test (SACT) testing during (A) 10% ethanol, (C) water, and (E) 20% sucrose conditions in the full cohort (*n* = 15). (B, D, F) Zone occupancy during the corresponding (B) ethanol, (D) water, and (F) sucrose conditions in baseline-defined low-CSI (*n* = 7) and high-CSI (*n* = 8) mice. (G) Post-CIE Chamber Sociability Index (CSI) across bottle conditions in low- and high-CSI mice. (H, I) Paired baseline-to-post-CIE CSI values in (H) low-CSI and (I) high-CSI mice. (J) Distance traveled and (K) zone transitions during post-CIE SACT testing across bottle conditions. Individual animals are shown; bars represent mean ± SEM, and violin plots show individual distributions with median and quartiles. Zone occupancy was analyzed separately by bottle condition using two-way repeated-measures ANOVA. Whole-cohort baseline-to-post-CIE comparisons were followed by Fisher’s LSD for the ethanol condition and Šídák-adjusted comparisons for the water and sucrose conditions; exploratory CSI-stratified comparisons were followed by Tukey-adjusted comparisons for ethanol and sucrose and Šídák-adjusted comparisons for water. Post-CIE CSI was analyzed by two-way repeated-measures ANOVA with bottle condition as a within-subject factor and CSI stratum as a between-subject factor, followed by Tukey-adjusted comparisons. Longitudinal CSI was analyzed separately within each CSI stratum by two-way repeated-measures ANOVA with bottle condition and session as within-subject factors, followed by Tukey-adjusted comparisons. Distance traveled and zone transitions were analyzed by one-way repeated-measures ANOVA with Tukey-adjusted comparisons. *P* < 0.05, P < 0.01, *P* < 0.001, P < 0.0001. CIE, chronic intermittent ethanol; SACT, Social–Alcohol Choice Test; CSI, Chamber Sociability Index.

During the water condition, the whole-cohort zone × session interaction was significant (*F*(1, 14) = 49.52, *P* < 0.0001). Šídák-adjusted comparisons showed increased Social Zone time from 475.3 to 592.7 s (adjusted *P* < 0.0001) and decreased Bottle Zone time from 278.1 to 206.1 s (adjusted *P* = 0.0040; Figure 4C). Paired longitudinal comparisons within each baseline CSI stratum showed the same directional pattern. In low-CSI mice, Social Zone time increased from 460.1 to 572.5 s (adjusted *P* = 0.0005), whereas Bottle Zone time decreased from 287.5 to 205.8 s (adjusted *P* = 0.0080). In high-CSI mice, Social Zone time increased from 488.5 to 610.3 s (adjusted *P* = 0.0055), and Bottle Zone time decreased from 270.0 to 206.3 s (adjusted *P* = 0.0420; Figure 4D).

During the 20% sucrose condition, neither the whole-cohort zone × session interaction (*F*(1, 14) = 0.02716, *P* = 0.8715) nor the main effect of session (*F*(1, 14) = 2.155, *P* = 0.1642) was significant. Šídák-adjusted comparisons likewise detected no baseline-to-post-exposure change in Social Zone or Bottle Zone time (adjusted *P* = 0.9370 and 0.9947, respectively; Figure 4E). Consistent with the whole-cohort analysis, paired comparisons within the fixed CSI strata detected no baseline-to-post-exposure changes in Social Zone time in low-CSI or high-CSI mice (adjusted *P* = 0.9393 and 0.5454, respectively) or in Bottle Zone time (adjusted *P* = 0.9812 and 0.9954, respectively; Figure 4F).

At post-exposure SACT testing, CSI differed across the three bottle conditions (Geisser– Greenhouse-corrected main effect of condition, *F*(1.500, 19.50) = 25.87, *P* < 0.0001), whereas the main effect of baseline CSI stratum (*F*(1, 13) = 0.8921, *P* = 0.3621) and condition × CSI-stratum interaction (*F*(1.500, 19.50) = 0.7700, *P* = 0.4415) were not significant. Tukey-adjusted paired comparisons showed lower CSI during ethanol than water testing in both low-CSI and high-CSI mice (adjusted *P* < 0.0001 and 0.0008, respectively). In high-CSI mice, CSI was also lower during ethanol than sucrose testing (adjusted *P* = 0.0080). Low-CSI and high-CSI mice did not differ significantly from one another during ethanol, water, or sucrose testing (adjusted *P* = 0.8503, 0.1856, and 0.3547, respectively; Figure 4G).

Longitudinal CSI analyses retained the same individual animals across all three bottle conditions and both testing sessions. In low-CSI mice, there were significant main effects of session (*F*(1, 6) = 40.56, *P* = 0.0007) and bottle condition (*F*(1.088, 6.526) = 7.432, *P* = 0.0301), whereas the condition × session interaction did not reach significance (*F*(1.108, 6.648) = 5.538, *P* = 0.0506). Tukey-adjusted paired comparisons showed increased CSI from baseline to post-exposure during water testing (0.03851 to 0.2866; adjusted *P* = 0.0001), with no significant change during ethanol testing (−0.02601 to −0.03951; adjusted *P* = 0.7808) or sucrose testing (−0.1201 to 0.04316; adjusted *P* = 0.0553; Figure 4H).

In high-CSI mice, there was a significant main effect of bottle condition (*F*(1.740, 12.18) = 5.968, *P* = 0.0182) and a significant condition × session interaction (*F*(1.775, 12.43) = 26.73, *P* < 0.0001), whereas the main effect of session was not significant (*F*(1, 7) = 4.192, *P* = 0.0798). Tukey-adjusted paired comparisons showed that CSI decreased from 0.1516 at baseline to −0.08513 post-exposure during ethanol testing (adjusted *P* = 0.0045), whereas CSI increased from 0.09956 to 0.3646 during water testing (adjusted *P* = 0.0024) and from −0.06692 to 0.1534 during sucrose testing (adjusted *P* = 0.0109; Figure 4I).

Supporting analyses incorporating all three chamber zones at post-exposure SACT testing further characterized condition-dependent spatial allocation. In low-CSI mice, the zone × bottle-condition interaction was significant (*F*(1.620, 9.723) = 5.765, *P* = 0.0266). Social Zone time was lower and Bottle Zone time higher during ethanol than water testing (adjusted *P* = 0.0002 and 0.0095, respectively), whereas Neutral Zone comparisons were not significant (Figure S1C). In high-CSI mice, the zone × bottle-condition interaction was also significant (*F*(1.760, 12.32) = 25.83, *P* < 0.0001). Social Zone time was lower during ethanol than water or sucrose testing (adjusted *P* < 0.0001 and 0.0027, respectively), whereas Neutral and Bottle Zone times were higher during ethanol than water (adjusted *P* = 0.0114 and 0.0018, respectively) and sucrose (adjusted *P* = 0.0190 and 0.0439, respectively; Figure S1D).

Post-exposure locomotor measures were also compared across bottle conditions using repeated measurements from the same 15 mice. Distance traveled during the 15-minute SACT differed significantly across ethanol, water, and sucrose conditions (Geisser–Greenhouse-corrected one-way repeated-measures ANOVA, *F*(1.846, 25.84) = 12.39, *P* = 0.0002). Tukey-adjusted paired comparisons showed less distance traveled during sucrose testing than during ethanol or water testing (adjusted *P* = 0.0081 and 0.0008, respectively), whereas ethanol and water did not differ significantly (adjusted *P* = 0.2383; Figure S2A). The omnibus effect of bottle condition on zone transitions was also significant (*F*(1.716, 24.03) = 4.762, *P* = 0.0222), although none of the Tukey-adjusted pairwise comparisons reached significance (ethanol vs water, adjusted *P* = 0.6699; ethanol vs sucrose, adjusted *P* = 0.0574; water vs sucrose, adjusted *P* = 0.0975; Figure S2B).

Together, the analyses in Figure 4 identified condition-specific longitudinal changes in zone occupancy and CSI following CIE exposure, first at the whole-cohort level and then within the exploratory baseline CSI strata. To determine how these group-level patterns were reflected in individual animals, we next examined the paired baseline-to-post-exposure trajectories of each mouse within the fixed low-CSI and high-CSI strata (Figure 5).

**Figure 5.**
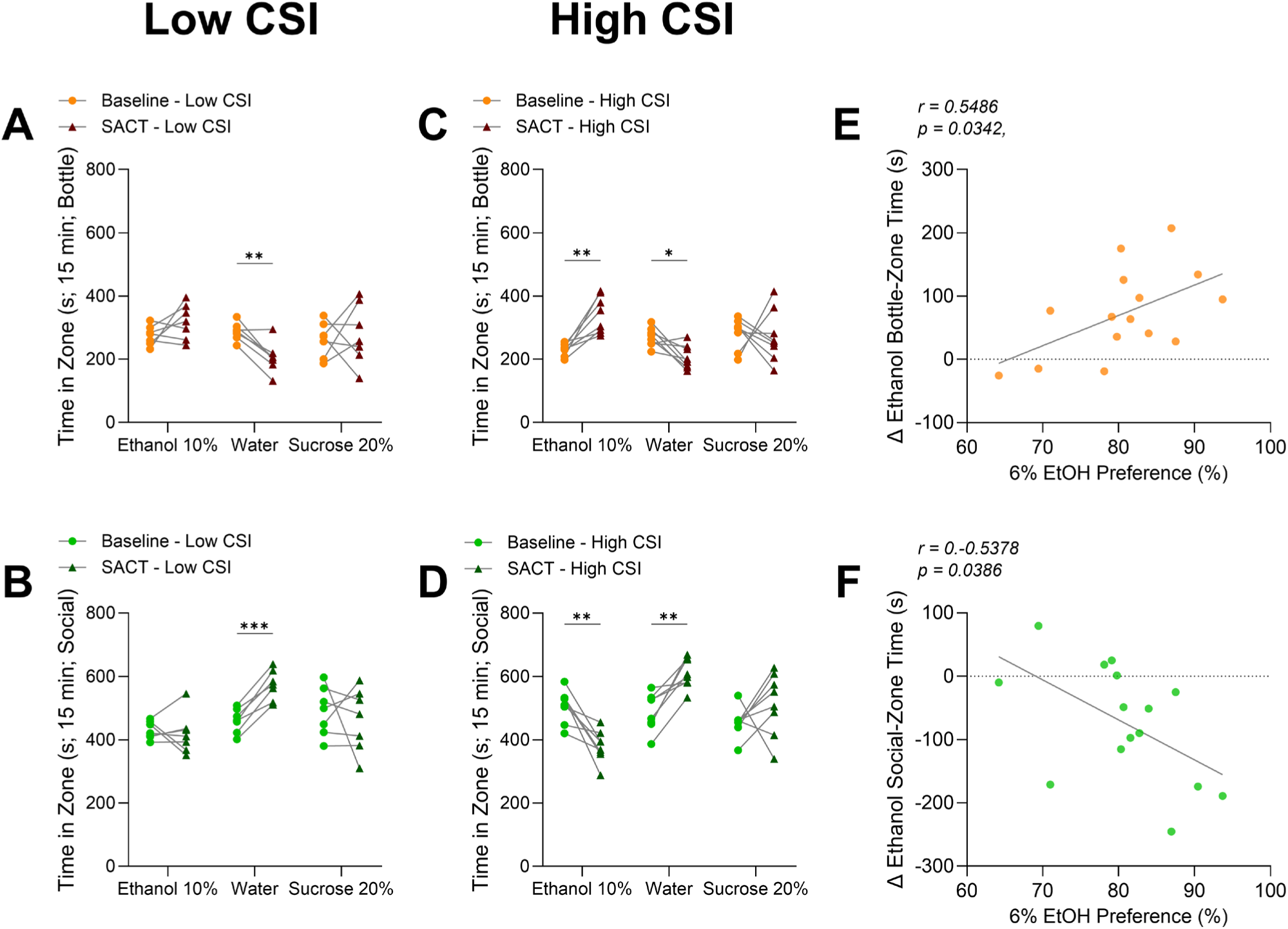
Individual longitudinal SACT trajectories within baseline-defined CSI strata and their association with ethanol preference. (A,B) Paired baseline-to-post-CIE Social–Alcohol Choice Test (SACT) trajectories in low-CSI mice (*n* = 7) for (A) Bottle Zone and (B) Social Zone time across 10% ethanol, water, and 20% sucrose conditions. (C,D) Corresponding paired trajectories in high-CSI mice (*n* = 8) for (C) Bottle Zone and (D) Social Zone time. Lines connect measurements from the same animal. (E,F) Association of 6% ethanol preference with baseline-to-post-CIE change (post-CIE minus baseline) in (E) ethanol-condition Bottle Zone time and (F) Social Zone time across all mice (*n* = 15). Panels A–D were analyzed separately within each CSI stratum by two-way repeated-measures ANOVA with bottle condition and session as within-subject factors, followed by Šídák-adjusted baseline-to-post-CIE comparisons within each bottle condition. Panels E and F were analyzed by simple linear regression, with significance determined by testing whether the regression slope differed from zero. *P* < 0.05, P < 0.01, *P* < 0.001. CIE, chronic intermittent ethanol; SACT, Social–Alcohol Choice Test; CSI, Chamber Sociability Index; EtOH, ethanol.

### SACT captured individual longitudinal trajectories within exploratory baseline CSI strata

To visualize and quantify within-animal longitudinal change, the same mice were tracked from their own baseline measurements through post-exposure SACT testing within the fixed low-CSI (*n* = 7) and high-CSI (*n* = 8) strata (Figure 5). Whereas the preceding analyses summarize baseline and post-exposure differences at the cohort and stratum levels, Figure 5 emphasizes the paired trajectories of individual animals across the ethanol, water, and sucrose conditions. Bottle condition and session were therefore treated as repeated within-subject factors in these within-stratum analyses.

In low-CSI mice, Bottle Zone time showed a significant main effect of bottle condition (*F*(1.879, 11.28) = 4.132, *P* = 0.0470), but no main effect of session (*F*(1, 6) = 0.8198, *P* = 0.4001) or condition × session interaction (*F*(1.380, 8.280) = 2.974, *P* = 0.1157). Šídák-adjusted paired comparisons showed decreased Bottle Zone time from baseline to post-exposure during water testing (adjusted *P* = 0.0080), whereas changes during ethanol and sucrose testing were not significant (adjusted *P* = 0.3966 and 0.9812, respectively; Figure 5A). Because the condition × session interaction was not significant, these simple comparisons were not interpreted as evidence that the magnitude of longitudinal change differed across bottle conditions. Social Zone time in low-CSI mice showed a significant condition × session interaction (*F*(1.101, 6.604) = 5.736, *P* = 0.0479). Social Zone time increased from baseline to post-exposure during water testing (adjusted *P* = 0.0005), with no significant change during ethanol or sucrose testing (adjusted *P* = 0.9674 and 0.9393, respectively; Figure 5B).

In high-CSI mice, Bottle Zone time showed a significant condition × session interaction (*F*(1.354, 9.480) = 12.97, *P* = 0.0034). Šídák-adjusted paired comparisons showed increased Bottle Zone time during ethanol testing, from 229.4 to 339.8 s (adjusted *P* = 0.0047), and decreased Bottle Zone time during water testing, from 270.0 to 206.3 s (adjusted *P* = 0.0420), whereas Bottle Zone time did not change significantly during sucrose testing (adjusted *P* = 0.9954; Figure 5C). Social Zone time also showed a significant condition × session interaction (*F*(1.399, 9.794) = 32.99, *P* < 0.0001). Social Zone time decreased during ethanol testing, from 507.6 to 380.1 s (adjusted *P* = 0.0080), and increased during water testing, from 488.5 to 610.3 s (adjusted *P* = 0.0055), with no significant change during sucrose testing (adjusted *P* = 0.5454; Figure 5D). These within-stratum analyses characterize paired longitudinal trajectories within each exploratory baseline CSI stratum and do not constitute a direct statistical test of whether the magnitude of change differed between low-CSI and high-CSI mice.

Neutral Zone trajectories provided additional information on individual longitudinal spatial allocation. In low-CSI mice, the condition × session interaction was significant (*F*(1.375, 8.253) = 6.027, *P* = 0.0317), although Šídák-adjusted paired baseline-to-post-exposure comparisons were not significant during ethanol, water, or sucrose testing (adjusted *P* = 0.3510, 0.5753, and 0.4633, respectively; Figure S1E). In high-CSI mice, Neutral Zone time showed a significant main effect of session (*F*(1, 7) = 10.75, *P* = 0.0135), decreasing from a mean of 138.4 s at baseline to 103.0 s post-exposure when averaged across bottle conditions (mean difference, baseline minus post-exposure, 35.36 s; 95% CI, 9.865 to 60.86). The condition × session interaction was not significant (*F*(1.540, 10.78) = 1.221, *P* = 0.3196). Šídák-adjusted paired comparisons showed decreased Neutral Zone time during water testing, from 130.1 to 78.07 s (adjusted *P* = 0.0345), whereas changes during ethanol and sucrose testing were not significant (adjusted *P* = 0.7275 and 0.1571, respectively; Figure S1F). Because the condition × session interaction was not significant, the significant water simple comparison was not interpreted as evidence that the longitudinal change during water differed significantly from the changes observed during ethanol or sucrose testing.

### Ethanol preference was associated with individual longitudinal changes in ethanol-condition zone occupancy

The individual longitudinal changes quantified by the SACT were next examined in relation to voluntary ethanol preference independently of the exploratory CSI stratification. Across all 15 mice, 6% ethanol preference was positively associated with the individual baseline-to-post-exposure change in Bottle Zone time during ethanol testing (simple linear regression, *F*(1, 13) = 5.596, *P* = 0.0342, *R*² = 0.3009; slope, 4.804 s per percentage point; 95% CI, 0.4167 to 9.191; Figure 5E). Conversely, ethanol preference was negatively associated with the individual baseline-to-post-exposure change in Social Zone time during the ethanol condition (*F*(1, 13) = 5.291, *P* = 0.0386, *R*² = 0.2893; slope, −6.320 s per percentage point; 95% CI, −12.25 to −0.3843; Figure 5F). Thus, mice with greater voluntary ethanol preference tended to show a larger longitudinal shift in ethanol-condition spatial allocation toward the Bottle Zone and away from the Social Zone. These relationships indicate covariation between ethanol preference and individual longitudinal changes in SACT behavior and do not establish causation.

## Discussion

The Social Alcohol Choice Test (SACT) provided a longitudinal framework to examine how animals allocated time between a social stimulus and a bottle-associated option. Following three weeks of chronic intermittent ethanol (CIE) exposure, ethanol testing led to increased Bottle Zone occupancy and decreased Social Zone occupancy. In contrast, water testing increased Social Zone occupancy and decreased Bottle Zone occupancy, whereas sucrose testing did not significantly affect either measure. Notably, changes seen during ethanol sessions were correlated with independently measured ethanol preference. Together, these findings support further exploration of longitudinal spatial allocation as a complementary metric to traditional consumption-based measures.

The SACT extends established three-chamber social-approach procedures (Nadler et al., 2004, Yang et al., 2011) Moreover, incorporates the modified alcohol–social testing arrangement described by Su et al. (2023). Operant studies similarly demonstrate that alcohol and social access serve as competing alternatives, with allocation influenced by response requirements and experimental context (Marchant et al., 2023, Augier et al., 2023). Unlike discrete-choice procedures, which make outcomes mutually exclusive within a trial, the SACT measures continuous allocation across simultaneously accessible zones. Its key advantage is the integration of pre-exposure characterization, repeated testing of the same animals, and the inclusion of ethanol, water, and sucrose comparator conditions.

The observed trajectories underscore the importance of accounting for available alternatives when interpreting alcohol-related behavior. Behavioral-economic models and alcohol-versus-nondrug choice paradigms consistently show that behavioral allocation depends on the nature and salience of the competing options (Bickel et al., 2014, Augier et al., 2018). While repeated ethanol vapor exposure and withdrawal can increase subsequent voluntary drinking (Becker and Lopez, 2004)Colarusso and Weiner (2026) reported reduced ethanol-directed choice and intake during chronic vapor exposure when sweetened ethanol and sucrose were concurrently available. These findings provide context for the present results, but they do not establish shared mechanisms across species, exposure schedules, or endpoints. In this study, comparator sessions served as reference contexts rather than as exposure controls, and they did not consistently show a loss of Social Zone occupancy or an increase in Bottle Zone occupancy. Furthermore, separate within-condition analyses do not directly demonstrate that longitudinal changes differed statistically across bottle conditions, and nonsignificant sucrose changes do not confirm that reward valuation was preserved or that equivalence between sessions was maintained.

Exploratory stratification by Chamber Sociability Index (CSI) provided a descriptive assessment of individual variation. Among high-CSI mice, ethanol testing decreased Social Zone and increased Bottle Zone occupancy, whereas similar changes were not observed in low-CSI mice. Both groups increased Social Zone occupancy during water testing. These findings reflect changes relative to each animal’s baseline; however, statistical significance in one stratum but not another does not indicate differential responsiveness between strata (Nieuwenhuis et al., 2011). Post-exposure CSI analysis showed neither a significant main effect of stratum nor a condition-by-stratum interaction. Maintaining the identities of the same 7 low-CSI and 8 high-CSI mice across sessions, as illustrated in the paired trajectories in Figure 5, demonstrates this heterogeneity without treating the two time points as independent samples. Thus, the strata characterize longitudinal behavior rather than establish differential vulnerability to CIE.

Longitudinal allocation is also related to conventional drinking behavior. Although ethanol consumption and preference did not differ significantly between CSI strata, higher 6% ethanol preference correlated with greater increases in ethanol Bottle Zone time and greater decreases in Social Zone time across all 15 mice. Separate regression analyses accounted for approximately 30% of the variance in each change measure. Importantly, preference was measured during a vapor-free interval before the final SACT testing, rather than as an outcome following the behavioral assay. Consequently, these associations do not establish prospective prediction of drinking or demonstrate that preference caused the spatial changes. Instead, they indicate that within-animal allocation changes contain information related to voluntary ethanol preference and warrant independent replication.

The broader literature advises caution in interpreting sociability as uniformly protective against alcohol use. Human studies link extraversion to heavier alcohol consumption (Hakulinen et al., 2015)and experimental research demonstrates alcohol-related enhancement of social bonding and greater mood enhancement among more extraverted individuals in social contexts (Sayette et al., 2012, Fairbairn et al., 2015). Social and enhancement motives further contextualize associations between extraversion and drinking (Hussong, 2003, Kuntsche et al., 2005). Preclinical findings also depend on biological context: CIE withdrawal reduced sociability in DBA/2J but not in C57BL/6J males under the conditions examined by Sidhu et al. (2018), and a multivariable behavioral profile, including sociability, predicted ethanol preference in adolescent but not adult mice (Portugal et al., 2026). These findings underscore the importance of studying behavioral heterogeneity without equating mouse CSI with human extraversion or presuming that baseline social allocation universally predicts drinking.

The behavioral endpoints also require careful interpretation. While social experience can be rewarding in mice (Dölen et al., 2013)chamber occupancy does not equate to direct investigation, sustained affiliative interaction, or reinforcement (Yang et al., 2011, Sofyana et al., 2026). The CSI summarizes Social Zone occupancy relative to the combined Neutral and Bottle Zones and is not a direct measure of social-versus-ethanol preference. A negative CSI does not necessarily indicate greater Bottle Zone than Social Zone occupancy; examining individual zones is essential. Similarly, Bottle Zone time alone does not confirm drinking or alcohol seeking. Incorporating lickometer, session-specific fluid measurements, stimulus-cage proximity, and direct social-investigation measures would clarify which behaviors contribute to changes in spatial allocation.

Several design limitations constrain the interpretation of these findings. Because all animals underwent CIE, the absence of a longitudinal air-exposed control group prevents disentangling exposure-related changes from the effects of repeated testing, aging, handling, or other time-dependent variables. The current design also does not separate the specific consequences of chronic ethanol exposure from those attributable to withdrawal. Water and sucrose sessions served only as within-subject comparators, not as true exposure controls. Baseline testing involved two sessions per condition, whereas post-exposure testing involved only one, and both phases used a fixed sequence of ethanol, water, and sucrose, with post-exposure testing starting 24 hours after the final vapor exposure. Consequently, condition was confounded with testing day, order, and withdrawal interval, and stimulus locations were not fully counterbalanced between phases. Employing matched sampling schedules and counterbalanced testing would strengthen interpretability.

The small cohort of male C57BL/6J mice further limits generalizability and precision, especially following stratification. Since CSI strata were defined based on baseline allocation, their initial differences do not independently validate distinct sociability phenotypes, and regression to the mean may influence subsequent changes {Barnett, 2005 #243}. Nonsignificant comparisons should not be interpreted as evidence of equivalence. Larger studies should assess CSI as a continuous variable, evaluate baseline stability, and replicate associations with preferences across sexes and diverse biological backgrounds. Activity-related factors also remain unresolved: open-field locomotion decreased, and post-exposure SACT distance varied across conditions. Because open-field testing occurred before final SACT testing, session-matched activity measures are necessary to distinguish altered allocation from changes in exploration or movement.

In conclusion, the SACT demonstrates proof of concept for longitudinal assessment of social-versus-bottle spatial allocation. Monitoring the same animals across baseline and post-exposure testing revealed individual trajectories of ethanol preference, measured independently. These findings position the assay as a complement to established measures of consumption and reinforcement. To establish its reproducibility, exposure specificity, and broader utility, future studies should incorporate exposure controls, matched testing schedules, larger and more diverse cohorts, and direct measures of drinking and social investigation.

## Ethics and Integrity Statement

The authors confirm that this manuscript represents original work, has not been published previously except as permitted by *Alcohol: Clinical and Experimental Research (ACER)*, and is not currently under consideration for publication elsewhere. All authors have approved the submitted version of the manuscript and agree to its submission to *ACER*. The authors affirm that the work complies with the journal’s applicable ethical, legal, and research integrity requirements.

## Data Availability Statement

The data generated and analyzed during the current study are available from the corresponding author upon reasonable request and will be deposited in an appropriate public repository in accordance with the data-sharing requirements of *Alcohol: Clinical and Experimental Research*.

## Author Contributions

**Nathan Jones:** Conceptualization, Methodology, Investigation, Formal Analysis, Data Curation, Visualization, Writing – Original Draft, Writing – Review & Editing. **Elliot Arozqueta:** Conceptualization, Methodology, Investigation, Formal Analysis, Data Curation, Visualizatio**n. Owen Alt:** Methodology, Investigation, Formal Analysis. **Hannah Aranda:** Investigation, Formal Analysis. **Phillip Starski:** Formal Analysis, Visualization, Writing – Review & Editing. **Doo-Sup Choi:** Writing – Review & Editing, Supervision.

## Ethics Approval Statement

All procedures involving mice were reviewed and approved by the Institutional Animal Care and Use Committee (IACUC) at Mayo Clinic under protocol A5370. All animal procedures were conducted in accordance with applicable institutional guidelines and regulations for the care and use of laboratory animals.

## Patient Consent Statement

Not Applicable

## Clinical Trial Registration

Not applicable.

## Acknowledgments and Disclosures

We thank all laboratory members for their helpful discussions and comments. Figures were created using BioRender.com. Mayo Clinic’s ChatGPT Enterprise (OpenAI) was used solely for editorial assistance to tighten and condense sentences after the authors had developed the manuscript and scientific content. The authors reviewed and approved all resulting revisions and take full responsibility for the manuscript’s content.

