## Supplemental Information for "Longitudinal Changes in Social–Alcohol Behavioral Allocation During Chronic Ethanol Exposure"

**Supporting Information for:** *Longitudinal Changes in Social–Alcohol Behavioral Allocation During Chronic Ethanol Exposure*

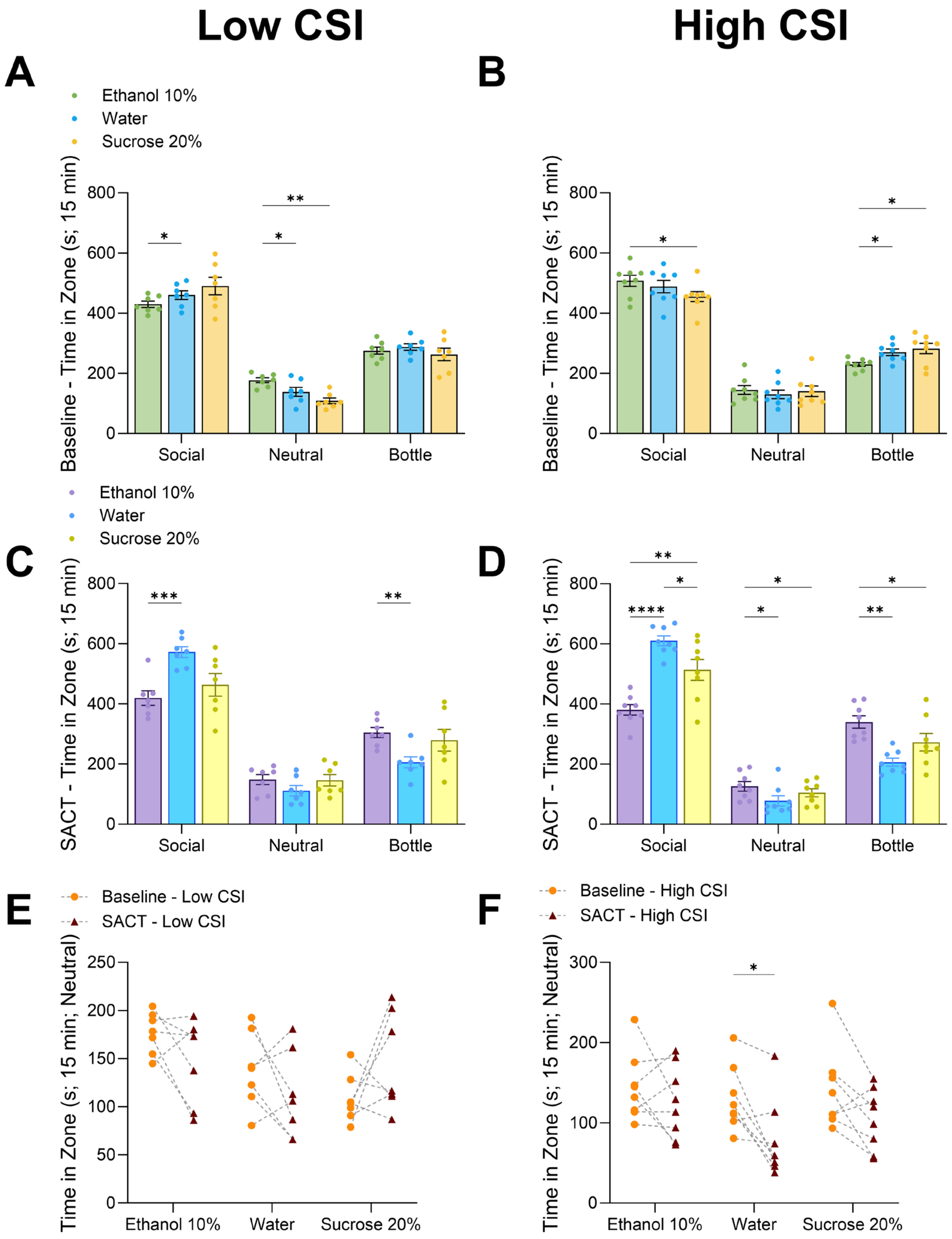

**Figure S1. Zone occupancy during baseline and post-exposure Social Alcohol Choice Test (SACT) according to baseline sociability phenotype.** Mice were stratified into low-CSI (*n* = 7) and high-CSI (*n* = 8) groups based on baseline CSI scores. **(A, B)** Baseline time spent in the social, neutral, and bottle zones during the 10% ethanol, water, and 20% sucrose conditions for low-CSI (A) and high-CSI (B) mice. **(C, D)** Post-exposure SACT time spent in the social, neutral, and bottle zones during the same solution conditions for low-CSI (C) and high-CSI (D) mice. Bars show group means with individual mice overlaid. **(E, F)** Within-subject changes in neutral-zone time from baseline to post-exposure SACT for low-CSI (E) and high-CSI (F) mice under each solution condition. Circles represent baseline measurements, triangles represent post-exposure SACT measurements, and dashed lines connect measurements from the same mouse. Zone occupancy is expressed as time in seconds during the 15-min test. Significant comparisons are indicated by brackets: * $P < 0.05$, ** $P < 0.01$, *** $P < 0.001$, and **** $P < 0.0001$.

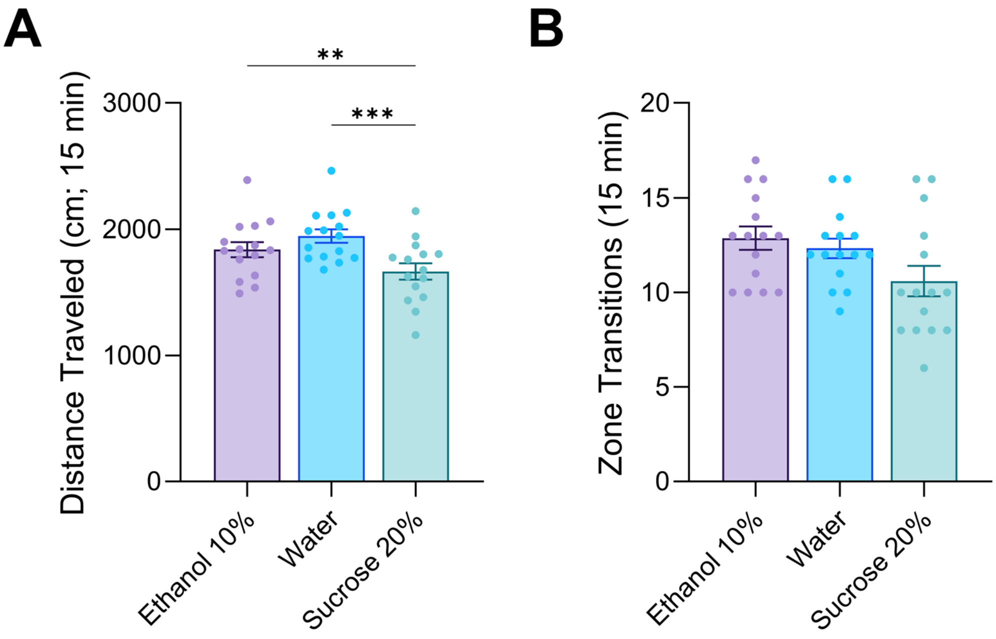


**Figure S2. Locomotor activity and zone transitions during the Social Alcohol Choice Test (SACT). (A)** Total distance traveled during the 15-min SACT under the 10% ethanol, water, and 20% sucrose conditions. Distance traveled differed across conditions (repeated-measures one-way ANOVA with Geisser-Greenhouse correction, $P = 0.0002$). Mice traveled a greater distance during the 10% ethanol condition than during the 20% sucrose condition ($P = 0.0081$) and during the water condition than during the 20% sucrose condition ($P = 0.0008$). **(B)** Number of zone transitions during the 15-min SACT under each solution condition. Zone transitions differed across conditions (repeated-measures one-way ANOVA with Geisser-Greenhouse correction, $P = 0.0222$), although no individual pairwise comparison reached statistical significance. Bars show group means with individual mice overlaid ($n = 15$). Error bars represent SEM. Post hoc comparisons were performed using Tukey's multiple-comparisons test. Significant pairwise comparisons are indicated by brackets: **P** < 0.01 and ***P*** < 0.001.
